# Brief Investigation: Investigating the role of phosphodiesterase Pde2 in coordinating the yeast Environmental Stress Response

**DOI:** 10.64898/2026.05.20.726645

**Authors:** Rachel A. Kocik, Jamie Ahrens, Audrey P. Gasch

**Affiliations:** Center for Genomic Science Innovation, University of Wisconsin-Madison, Madison, WI 53706; Department of Medical Genetics, University of Wisconsin-Madison, Madison, WI 53706

## Abstract

Yeast responding to acute stress reallocate cellular resources, in part via the Environmental Stress Response (ESR) that induces stress-defense genes while repressing ribosome-biogenesis and growth genes. The purpose and regulation of coordinated induction and repression is incompletely understood, but both responses are influenced by ESR transcription factors Msn2 and Msn4 (Msn2/4). Here we used single-cell microscopy and transcriptomic analysis to investigate the role of upstream regulator Pde2 in ESR regulation and post-stress fitness. Loss of *PDE2* weakened and shortened Msn2 activation following salt stress and produced muted induction of Msn2/4 targets, similar to a *msn2∆msn4∆* strain. In contrast, Pde2 had at most a minor impact on ESR repressor Dot6, yet was important for repression of its targets beyond Msn2/4 influence. Consistent with our recent resource-reallocation model, *pde2∆* cells had normal or faster post-stress growth rates, despite weaker activation of the ESR. We discuss implications for ESR regulation and function.

## INTRODUCTION

Mounting a rapid cellular response to stressful insults often requires re-allocation of cellular resources away from growth and division and toward cellular defense. While organisms have evolved in regulatory strategies, a common theme is the transcriptional and/or translational up-regulation of defense genes and proteins and the down-regulation of others that compete for transcriptional and translational machinery (Ho & Gasch, 2015; Kocik et al., 2026). We previously hypothesized that fast-growing budding yeast, *Saccharomyces cerevisiae*, accomplishes this reallocation in part through a common stress-activated transcriptional program called the Environmental Stress Response (ESR)(Gasch et al., 2000). Upon acute environmental stress, cells coordinately induce ∼300 genes involved in various aspects of stress defense (iESR genes) while repressing ∼600 genes encoding ribosomal proteins (RP), ribosome biogenesis factors (RiBi), and other proteins required for rapid growth (collectively, rESR genes).

Coupling iESR induction with rESR repression may aid in resource reallocation. Cells lacking RiBi repressors *DOT6* and paralog *TOD6* fail to properly repress RiBi genes during stress (Ho et al., 2018; Lippman & Broach, 2009). Instead, overly-abundant RiBi transcripts remain associated with ribosomes at the expense of iESR mRNAs that show delayed ribosome association and translation (Ho et al., 2018). iESR induction is not required to survive single-stress treatments, but it is critical for increased tolerance to subsequent stress exposures (Berry et al., 2011; Berry & Gasch, 2008; Guan et al., 2012). Consequently, delayed production of iESR proteins in *dot6∆tod6∆* cells results in delayed acquisition of secondary-stress tolerance (Kocik et al., 2026). But mounting the response comes at a cost: cells lacking iESR inducers Msn2 and Msn4 (Msn2/4) fail to properly mount the iESR during stress, but in fact grow *faster* than wild type after single-stress treatments. In contrast, cells lacking Dot6/Tod6 induce the iESR without concomitant RiBi repression, and grow slower than wild type after single-stress treatments (Bergen et al., 2022; Ho et al., 2018; Kocik et al., 2026). We proposed that rESR repression helps to balance the cost of iESR induction. Consist with this model, a quadruple mutant lacking all four regulators grows equal to or slightly faster than wild-type cells responding to stress.

Here we extended our recent work by investigating upstream regulator Pde2, which inhibits the Protein Kinase A (PKA) pathway implicated in both iESR and rESR regulation. Under optimal conditions, PKA directly phosphorylates Msn2/4 and Dot6/Tod6 to restrict them to the cytosol (Beck & Hall, 1999; Deminoff et al., 2006; Garreau et al., 2000; Huber et al., 2011; Lippman & Broach, 2009). Upon stress, PKA inhibition, along with positive effects of stress-activated pathways, promotes nuclear translocation of Msn2/4 and Dot6/Tod6 and transcriptional regulation of their targets. How PKA is suppressed during stress remains incompletely resolved, but one mechanism is through allosteric regulator cAMP. High cAMP levels activate PKA, by disrupting PKA association with inhibitory subunit Bcy1 (Kocik & Gasch, 2022). cAMP turnover is regulated by phosphodiesterases, including high-affinity Pde2 activated by stress (Hu et al., 2010; Ma et al., 1999; Nikawa et al., 1987). Pde2 physically interacts with several PKA-regulated transcription factors including Msn2, although the functional importance of that interaction is unknown (MacGilvray et al., 2018). Furthermore, how Pde2 affects Dot6 has not been studied previously. Here we investigated Msn2 and Dot6 activation dynamics, iESR and rESR transcriptome changes, and post-stress fitness in *pde2∆* cells responding to salt and other stresses. These results shed light on the regulatory network that influences ESR activation and expands on our perspective that ESR activation is not required to survive single-stress conditions.

## RESULTS AND DISCUSSION

### Pde2 affects Msn2 activation dynamics

We previously developed a microfluidics system to follow nuclear translocation dynamics of genomically encoded Msn2-mCherry and Dot6-GFP expressed in the same cells (Bergen et al., 2022; Kocik et al., 2026). Here, we investigated the factors’ activation dynamics in a *pde2∆* strain carrying both reporters. We mixed the mutant with a comparable wild-type strain containing a distinguishing iRFP marker, allowing mutant and wild type to be studied in the same microfluidics chamber. We measured Msn2-mCherry and Dot6-GFP nuclear translocation dynamics before and after exposure to an acute dose of 0.7M sodium chloride (NaCl) inducing osmotic and ionic stress (**Fig 1A**).

**Figure 1.**
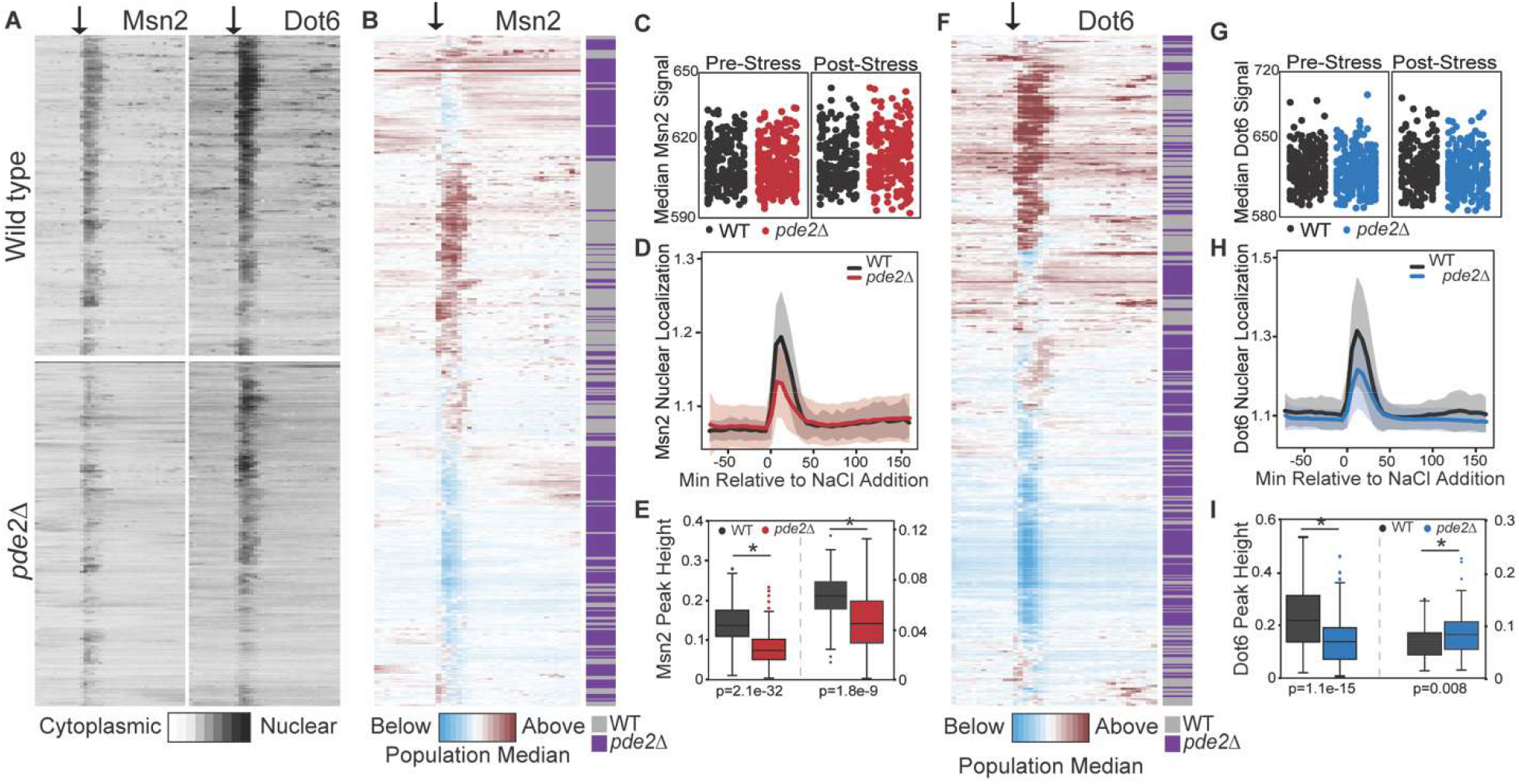
Pde2 regulates Msn2 nuclear translocation. Nuclear to cytoplasmic abundance of Msn2-mCherry or Dot6-GFP was measured for wild-type and *pde2∆* cells mixed in the same chamber. **A**) Nuclear/cytoplasmic ratio for 212 wild-type-iRFP and 256 *pde2∆* cells mixed in the same chamber over 5 biological replicates (strains shown separately for clarity). Each row represents a different cell and each column indicates a different time point spaced every 6 min. Time of NaCl addition is indicated by an arrow. **B and F**) log_2_ change in nuclear/cytoplasmic ratio for (**B**) Msn2-mCherry or (**F**) Dot6-GFP normalized to the median value of each population at each time point, according to the key. Cells were organized separately in each panel by hierarchical clustering, and cell genotype was subsequently annotated (purple, grey). Many *pde2∆* cells (purple) fell into separate clusters based on Msn2-mCherry patterns (B) consistent with weaker nuclear translocation. **C and G**) Distribution of protein abundances (median signal in each cell) before and after stress. **D and H**) Median trace of cells from (A), where shading represents one standard deviation. **E and I**) Distributions of nuclear-translocation peak heights as previously described (Kocik et al., 2026). The first pair of distributions on the left of each panel indicates when iRFP was expressed in wild-type cells; the second pair of distributions indicates when iRFP was expressed in *pde2∆* cells (117 wild type, 85 *pde2∆* cells). Signal intensity differs between the pairs (shown on left versus right axes) due to changes in lamp power over the span between these experiments. P-values, Wilcoxon rank-sum test comparing wild-type and *pde2∆* cells.

Cells lacking *PDE2* had a clear defect in Msn2 regulation after salt stress. Relative to wild type, *pde2∆* cells showed normal Msn2-mCherry abundance (**Fig 1B-C**) but significantly weaker nuclear translocation during NaCl stress (**Fig 1D-E**). The duration of nuclear Msn2-mCherry was also truncated, implicating Pde2 in maintaining nuclear Msn2 localization (**Fig 1D**). The defects in activation and nuclear retention were independent of which strain expressed the iRFP marker (**Fig 1E**), confirming that Pde2 influences both the magnitude and timing of Msn2 activation.

The effect of *pde2∆* on Dot6 was less clear. Past work showed that the *msn2∆msn4∆* mutant harbors less Dot6-GFP protein, we proposed in part through direct Msn2/4 binding of the *DOT6* promoter (Kocik et al., 2026). Here, *pde2∆* cells showed no difference in Dot6-GFP abundance (**Fig 1F-G**), suggesting that cells either harbor enough Msn2/4 activity to maintain Dot6-GFP or that Msn2/4 influence Dot6 abundance through a Pde2-independent mechanism. The effect on Dot6-GFP activity was not clear: although the peak height appeared lower when iRFP was expressed in wild-type cells (**Fig 1H**), the effect was inverted when iRFP was expressed in the mutant (**Fig 1I)**, due to an artifact of the iRFP marker (**Fig S1**). Thus, any Pde2-dependent effect on Dot6-GFP behavior is mild enough to be confounded by our system.

**Figure S1.**
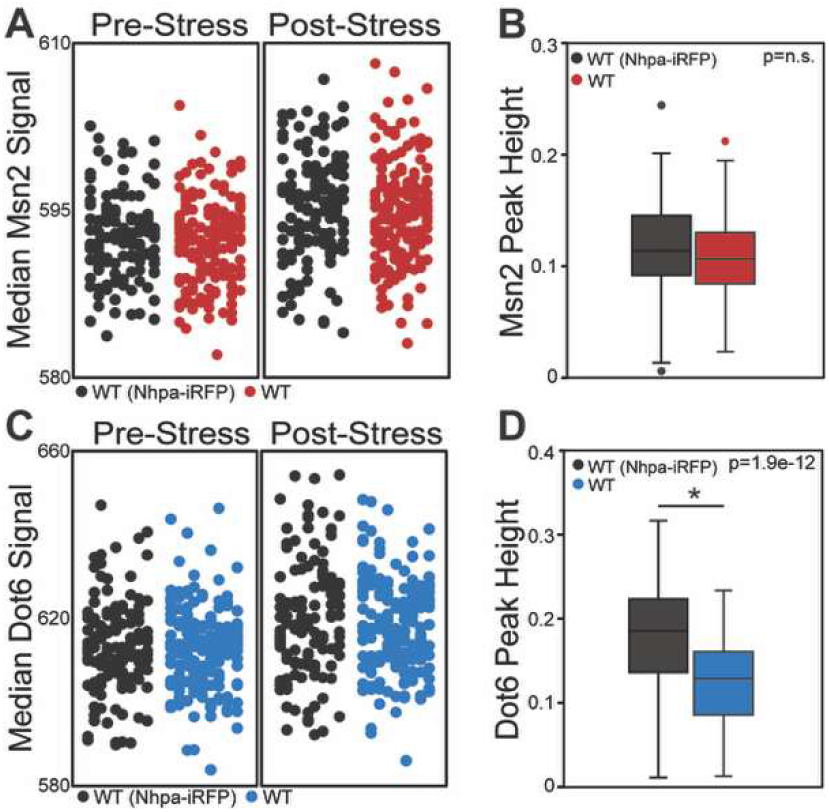
iRFP expression can affect peak height measurement in the GFP channel. Control experiments mixed two wild-type strains, both expressing Msn2-mCherry and Dot6-GFP but where only one strain carried the additional iRFP marker. **A and C**) Median pixel intensity of Msn2-mCherry (A) or Dot6-GFP (C) and **B and D**) nuclear translocation peak heights of Msn2-mCherry (B) and Dot6-GFP (D) are shown for the two strains. Cells carrying the iRFP marker reproducibly showed a larger apparent nuclear translocation signal in the GFP channel (see **D**), indicating an artifact of the iRFP marker. P-values based on Wilcoxon rank-sum test.

### Pde2 influences both iESR induction and rESR repression

We recently defined multiple sets of Msn2/4 targets based on temporal responses of wild-type, *msn2∆msn4∆*, and *dot6∆tod6∆* cells responding to 0.7M NaCl (Kocik et al., 2026). Here we investigated the genes’ salt-responsive dependence on Pde2 versus Msn2/4 using published transcriptomic data from our lab (Chasman et al., 2014). Induced genes in Clusters a-c (**Fig 2A**) showed very similar expression defects in *pde2∆* and *msn2∆msn4∆* cells studied in the same experiment (**Fig 2B**). All three clusters are enriched for iESR genes, and direct assessment of the 283 iESR genes confirmed similar dependencies on both sets of regulators (**Fig 2C**). Since *pde2∆* cells show a major defect in Msn2 activation, the simplest explanation is that the Pde2 effect is primarily through Msn2/4, although it is possible that other regulators that control subsets of iESR genes could also be influenced by Pde2. In contrast, repressed genes influenced by Msn2/4 showed greater dependence on Pde2 in side-by-side experiments (**Fig 2B**). This includes genes in Clusters d and e that are enriched for RiBi genes and Clusters f and g enriched for RP genes (Kocik et al., 2026). Furthermore, direct interrogation of those gene groups shows that the *pde2∆* defect is significantly greater than the *msn2∆msn4∆* defect (p < 8e-5, **Fig 2C**). This strongly suggests that Pde2 influences other regulators of rESR genes. One possibility is a direct connection to Dot6/Tod6: although we could not confirm a *pde2∆* defect in Dot6 activation due to fluorescent microscopy limitations, Dot6 can be regulated by PKA, raising the possibility of a direct connection (Deminoff et al., 2006; Kunkel et al., 2019; Lippman & Broach, 2009). However, we returned to previous data from our lab that identified proteins immunoprecipitated with Pde2, which included RP regulator Rap1 and RiBi regulator Abf1 (MacGilvray et al., 2018). The functional importance of these interactions, observed both before and after salt stress, is unknown; however, the interactions elevate these regulators as candidates through which Pde2 may impact rESR expression.

**Figure 2.**
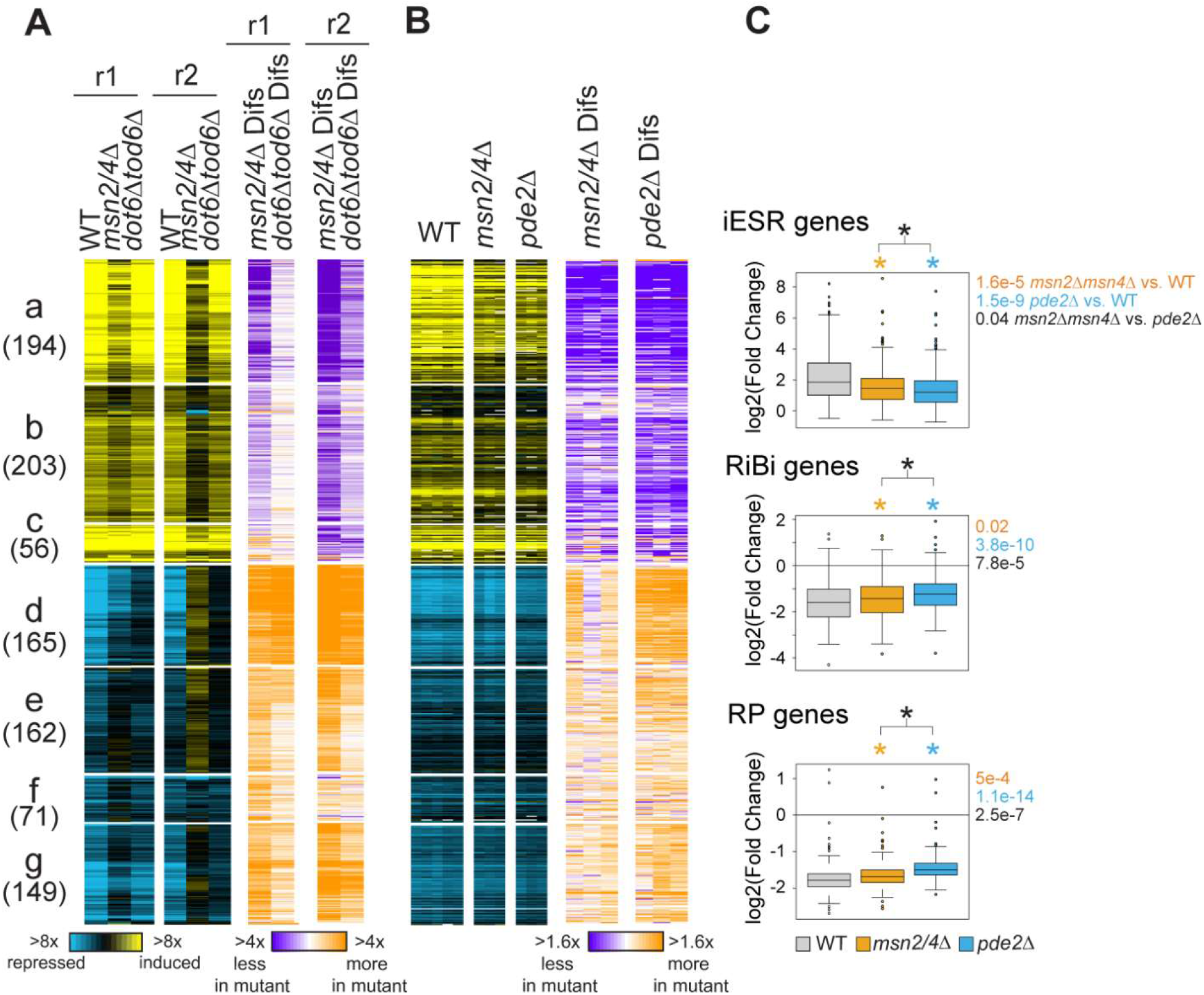
Contribution of Pde2 versus Msn2/4 to NaCl-responsive expression. Shown are 966 genes (rows) previously identified as induced or repressed in wild-type cells but with a defect in *msn2∆msn4∆* cells, as defined in (Kocik et al., 2026). The number of genes per cluster is indicated in parentheses. **A**) Log_2_(fold change) in expression in wild type, *msn2∆msn4∆, and dot6∆tod6∆* at 30 min after 0.7M NaCl compared to unstressed cells from replicate time courses (r1, r2), according to the key (Kocik et al., 2026). Differences between mutant and wild-type are shown according to the purple-orange key immediately below, for genes shown in A. **B**) As shown in A but for WT (n=5), *msn2∆msn4∆* (n = 3), and *pde2∆* (n = 3) from (Chasman et al., 2014). **C**) Distribution of log_2_(fold change) values for genes in the iESR (top), RiBi (middle), and RP (bottom) gene sets. Groups that were statistically different from wild type are indicated with an asterisk colored according to the key; groups different between *pde2∆* and *msn2∆msn4∆* are indicated in black text. Asterisk, p<0.05, Wilcoxon rank sum test.

### Cells lacking PDE2 grow normally or faster after stress treatments

Decoupling the ESR has significant effects on post-stress growth rate, but a quadruple mutant lacking *MSN2/MSN4/DOT6/TOD6* recovers with post-stress growth rates that are equal to or better than wild type (Kocik et al., 2026). Since *pde2∆* cells show a defect in both the iESR and rESR, we predicted post-stress growth rates close to wild-type cells. Indeed, in two of four stresses tested (ethanol and basic pH), *pde2∆* cells grew indistinguishably from wild type, whereas in response to low pH and salt studied here *pde2∆* cells grew slightly faster (**Fig 3**). We also assessed cells lacking the other low-affinity phosphodiesterase Pde1. Cells lacking both Pde2 and Pde1 grew worse after most stresses, and worse than either single-gene deletion, indicating that some cAMP phosphodiesterase activity is important for post-stress fitness.

**Figure 3.**
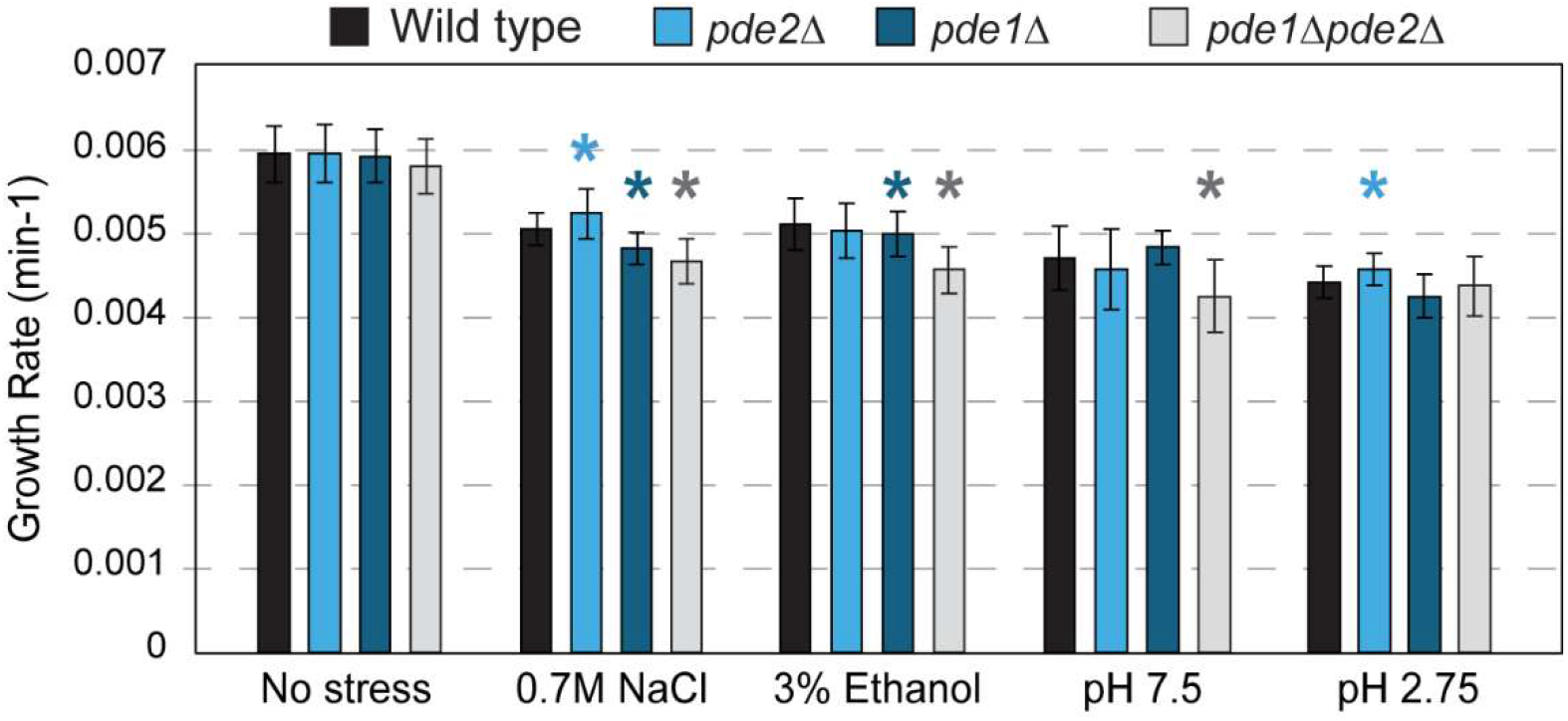
Pde2 is not required to maintain post-stress growth rate. Average and standard deviation of growth rates calculated for denoted strains growing in the absence of added stress (n = 12) or after addition of NaCl (n = 7), ethanol (n = 7), or high or low pH (n = 5). Asterisk indicates p < 0.05, replicated-paired T-tests.

### Concluding Remarks

Overall, this work expands understanding of ESR regulation and coordination, showing that Pde2 directly influences Msn2 activation dynamics and coordinates rESR repression through means beyond Msn2 input. This is consistent with the known inhibitory effect of PKA signaling on the ESR program (Garmendia-Torres et al., 2007; Klein & Struhl, 1994; Kocik & Gasch, 2022; Lippman & Broach, 2009; Smith et al., 1998). But several questions remain to be answered. How Pde2 is regulated during stress is not known, although it does undergo nucleocytoplasmic shuttling in a way that responds to PKA activity (Hu et al., 2010). Another question is if the physical interaction between Pde2 and PKA-regulated transcription factors, including Msn2, Abf1, and Rap1 discussed here, is important for transcriptional regulation. One possibility is that physical interactions and co-localized nuclear translocation is important for nuclear retention of Msn2, since the residual Msn2 nuclear localization in *pde2∆* cells appeared truncated in response to salt stress. Future studies will be required to dissect these possibilities.

## METHODS

### Strains and growth conditions

*Saccharomyces cerevisiae* strains of the BY4741 background used in this study are listed in **Table 1**. Strains were grown in Low Fluorescent Media (LFM) as previously described (0.17% yeast nitrogen base without ammonium sulfate, folic acid, or riboflavin; 0.5% ammonium sulfate; 0.2% complete amino acid supplement, and 2% glucose) (Bergen et al., 2022; Kocik et al., 2026). Strain AGY1577 (*pde1∆pde2∆*) was generated by replacing *PDE1* in the BY4741 *pde2::KANMX* strain (Open Biosystems) with the hygromycin-MX cassette via homologous recombination and validated using diagnostic PCRs. Crosses were used to generate AGY1575 (AGY1577 x AGY5), AGY1676 (AGY670 x AGY2228), and AGY2240 (AGY1676 x AGY2232). Liquid cultures for growth-rate assessment were grown as described in (Kocik et al., 2026). In brief, test tubes inoculated with overnight liquid cultures grown ∼12 hours in LFM were grown to an optical density at 600 nm (OD_600_) of ∼0.1 before measurements were taken every 15 minutes. Growth rates were calculated using an exponential fit of OD measurements during 75-225 min after addition of stress including: 0.7M NaCl or 3% ethanol added directly to the medium, or centrifugation and resuspension of cells in LFM media pH of either 7.5 or 2.75. Replicates were done on different days with paired wild-type and mutant cultures, enabling replicate-paired T-test analysis.

**Table 1.**
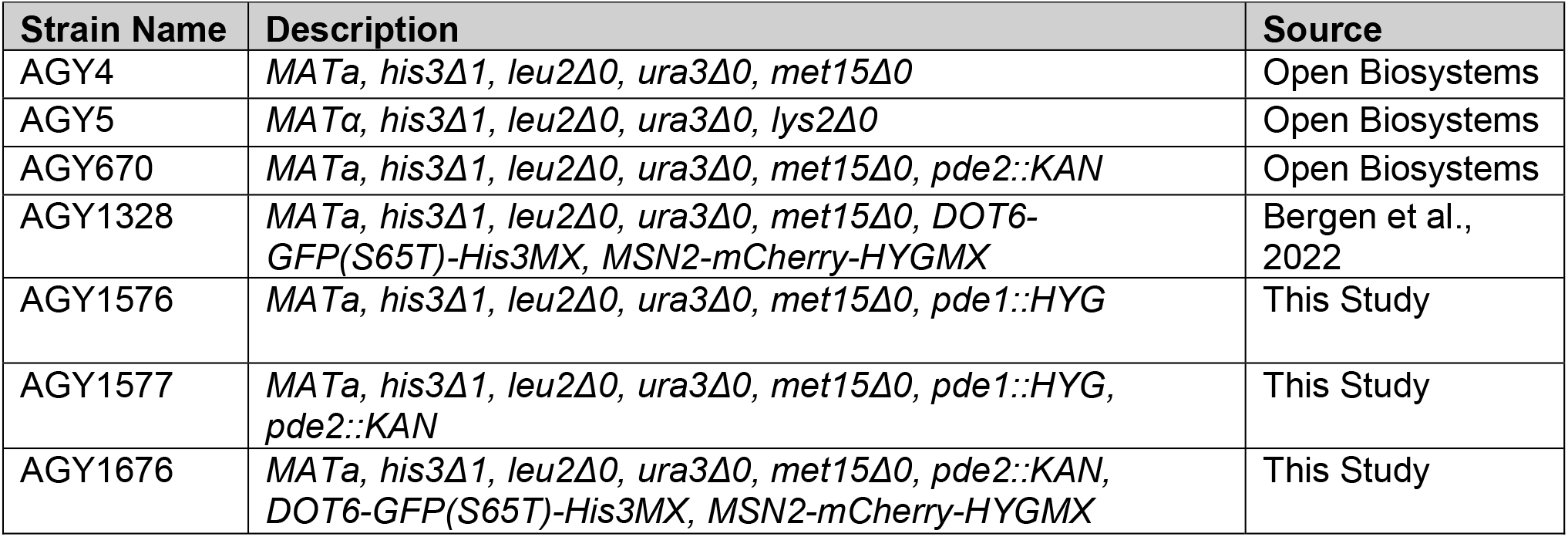
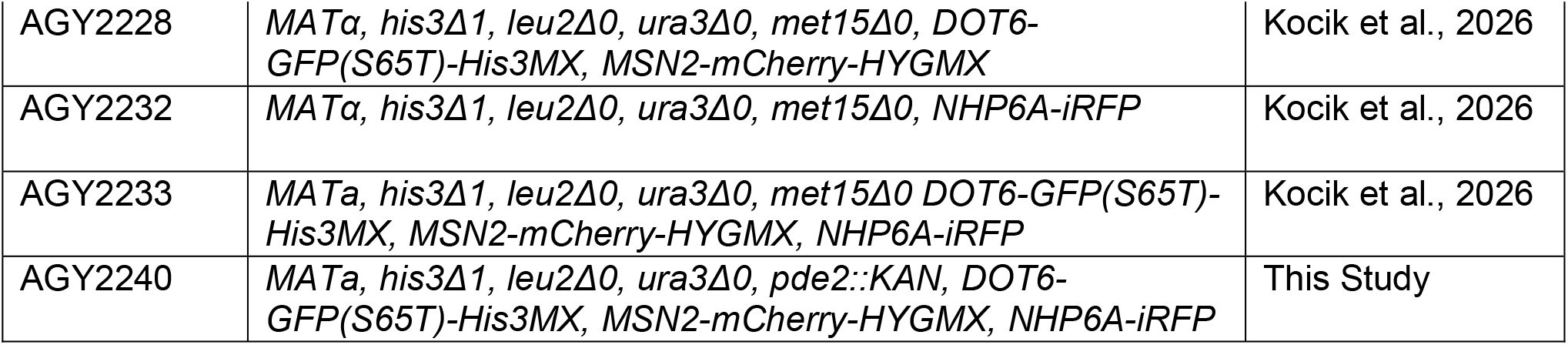
Strains used in this study.

### Microscopy and image analysis

Time-lapse microscopy was conducted using an FCS2 chamber (Bioptechs Inc, Butler, Pennsylvania) with image collection and analyses performed as previously described (Kocik et al., 2026). Msn2-mCherry and Dot6-GFP phenotypes were also determined as previously described, including “nuclear/cytoplasmic ratio” (defined as the average intensity of the top 5% of pixels divided by the median of all pixels, and acute stress peak height (defined as the maximum nuclear localization score immediately following NaCl addition (T13-T20) minus the minimum score just before NaCl was added (T11-T13)) (Bergen et al., 2022). Msn2 and Dot6 abundance were measured based on median Msn2-mCherry or Dot6-GFP signal in each cell at each timepoint. The average signal of each cell before (T1-T12) and after (T20-T36) NaCl addition was plotted in figures. Hierarchical clustering of cells in **Fig 1B** and **1F** was based on the log_2_(fold difference) compared to the population median. Clustering was performed using Gene Cluster 3.0 (Eisen et al., 1998) and visualized using Java TreeView version 1.2 (Saldanha, 2004).

### Transcriptomic analysis

Data from (Chasman et al., 2014) were analyzed in the context of gene sets identified in (Kocik et al., 2026) focusing on the subset of clusters with expression changes in the wild-type that are defective in *msn2∆msn4∆* cells. Transcription factor binding enriched for each cluster is available in (Kocik et al., 2026) Dataset EV2. The distribution of log_2_(fold change) values was taken for all ESR genes (Gasch et al., 2000). Distributions were compared using Wilcoxon rank-sum tests as indicated.

## DATA AVAILABILITY

Strains are available upon request. Dataset 1 (Kocik-pde2_Dataset-1_MicroscopyData.xlxs) contains all microscopy data. Dataset 2 (Kocik-pde2-Fig3-SourceData.xlxs) contains all growth curve data. Transcriptomic analysis used in this publication are from NIH GEO database and assigned the identifiers GSE283327 (Kocik et al., 2026) and GSE60613 (Chasman et al., 2014).

## ACKNOWLEDGEMENTS

We thank the McClean lab for reagents and microfluidics support (NIH grant R35GM128873 to Megan McClean), James Hose for help with RNA-sequencing, Michael Place for computational assistance, and Leah Escalante and members of the Gasch Lab for constructive discussions. This work was supported by NIH grant R01GM14975 to APG. RAK was supported by an NHGRI training grant to the Genomic Sciences Training Program T32HG002760.

## Supplemental Tables and Files

Dataset 1 (Kocik-pde2_Dataset-1_MicroscopyData.xlxs)

Dataset 2 (Kocik-pde2-Fig3-SourceData.xlxs)

